# Lef1 expression in fibroblasts maintains developmental potential in adult skin to regenerate wounds

**DOI:** 10.1101/2020.06.11.147066

**Authors:** Quan M. Phan, Gracelyn Fine, Lucia Salz, Gerardo G. Herrera, Ben Wildman, Iwona M. Driskell, Ryan R. Driskell

## Abstract

Scars are a serious health concern that impacts the clinical outcome and long-term well-being of burn victims and individuals with genetic skin conditions associated with wound healing. In this study using mouse as the model, we identify regenerative factors in neonatal skin that will transform adult skin to regenerate instead of repairing wounds with a scar, without perturbing normal development and homeostasis. We utilized single-cell RNA-sequencing (scRNA-seq) to probe unsorted cells from Regenerating, Scarring, Homeostatic, and Developing skin. Our results revealed a transient regenerative cell type in Developing skin, called papillary fibroblasts, which are defined by the expression of a canonical Wnt transcription factor Lef1. Tissue specific ablation of Lef1 inhibited skin regeneration. Importantly, ectopic expression of Lef1 in dermal fibroblasts did not disrupt development and aging, but primed adult skin to undergo enhanced regeneration. Here, we reveal the possibility of transferring the regenerative abilities of neonatal skin to adult tissue by expressing Lef1 in dermal fibroblasts. Finally, we have generated an expandable web resource with a search function to display gene expression in the context of our scRNA-seq data (https://skinregeneration.org/).

## Main

Understanding how to induce skin regeneration instead of scarring will have broad implications clinically and cosmetically (Walmsley et al., 2015b). One of the main characteristics of scars is the absence of hair follicles, indicating that their regeneration in a wound may be a critical step in achieving scar-less skin repair (Yang and Cotsarelis, 2010). Interestingly, human embryonic skin has the capacity to regenerate without scars (Lo et al., 2012). Similarly, neonatal and adult mouse skin has the capacity to regenerate small non-functional hair follicles under specific conditions (Figure 1c-d)(Ito et al., 2007; Rognoni et al., 2016; Telerman et al., 2017). These insights have prompted efforts in the field to define the molecular triggers that promote hair development in skin, with the ultimate goal of devising a way to regenerate fully functional hairs in adult skin wounds as a therapeutic modality (Yang and Cotsarelis, 2010).

**Figure 1:**
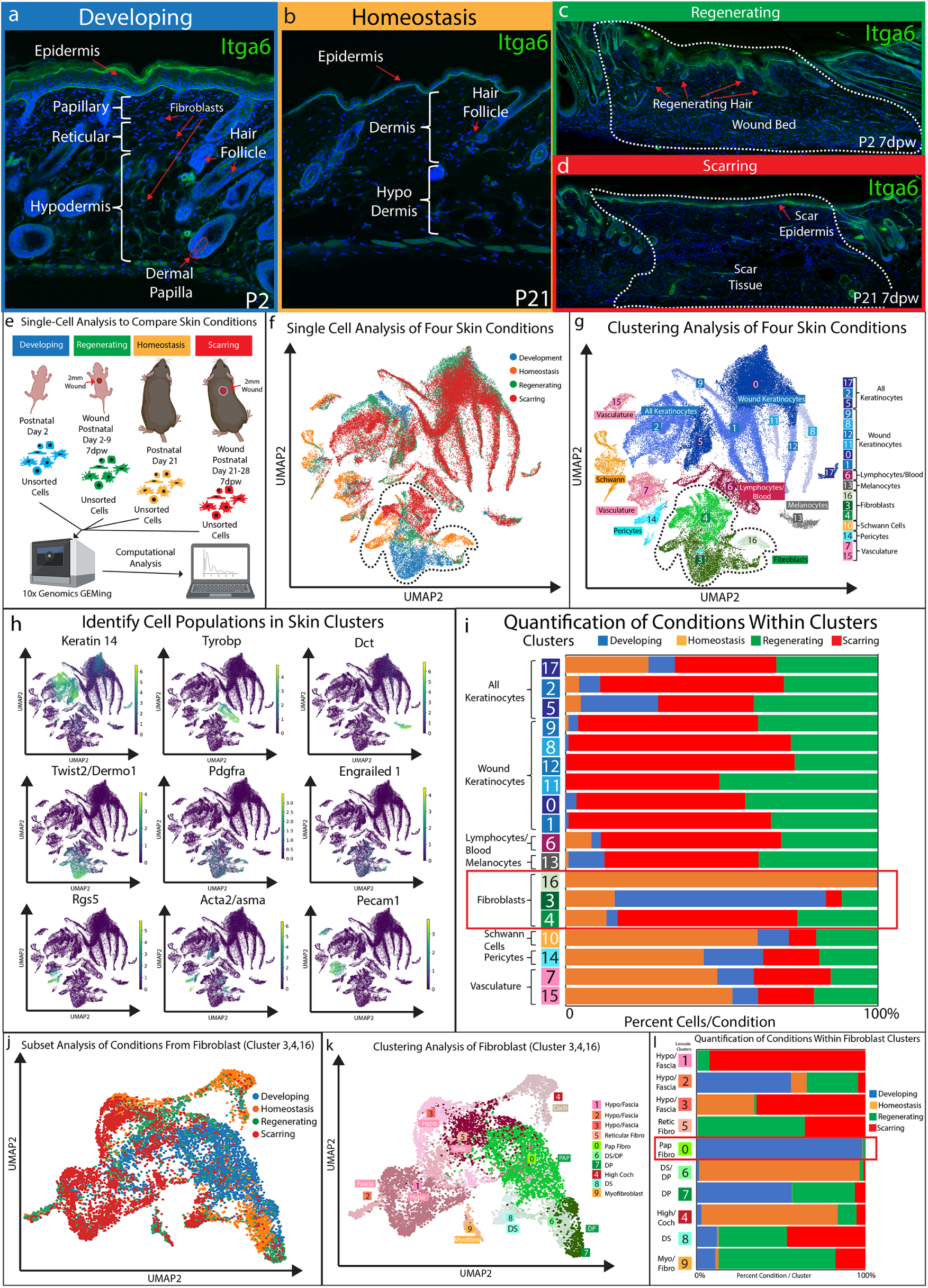
Fibroblasts in Developing skin are distinct from Regenerating, Scarring, and Homeostatic Conditions. (a-d) Immunostaining analysis and histological map of the dermis in Developing Post-natal day 2 (P2) skin (a), Homeostatic skin at P21 (b), skin undergoing regeneration P2 7 days post wound (7dwp) (c), and skin repairing a wound via scarring P21 7dpw (d). (e) Schematic representation of single cell isolation, library preparation, and sequencing analysis for unsorted cells from Developing (P2), Regenerating (P2 7dpw), Homeostatic (P21), and Scarring (P21 7dpw) conditions. (f) UMAP visualization of all cell populations for all conditions. Each cell is color coded based according to condition as labeled. (g) Clustering analysis of the UMAP plot, color coded based on cell types. (h) Overlaying gene expression on UMAP clusters to identify cell types. (i) Quantification of the percentage of all cells represented within a cluster. (j) Subset and re-clustering of Cluster 3,4,16 by computational integration. (k) Cluster analysis of integrated fibroblast clusters 3,4,16. (l) Quantitation of fibroblasts within each cluster represented by condition.

Human and mouse skin are similar in their overall structural complexity, indicating that mouse skin can be a useful model to study skin development and wound repair (Chen et al., 2013). Murine hair follicle and skin development primarily occurs between embryonic day 12.5 (E12.5) to post-natal day 21 (P21) (Muller-Rover et al., 2001). During this time different fibroblast lineages are established that respond to the changes in the environment to support hair follicle and skin development (Driskell et al., 2013; Jiang et al., 2018; Rinkevich et al., 2015; Rognoni et al., 2016). Fibroblasts that support hair follicle development differentiate from the papillary fibroblast lineage, into dermal papilla (DP), dermal sheath (DS), and arrector pili (AP) cells (Driskell et al., 2013; Plikus et al., 2017). Reticular fibroblasts, which cannot differentiate into papillary fibroblast lineages, secrete Extra-Cellular-Matrix and form into adipocytes (Driskell et al., 2013; Schmidt and Horsley, 2013). By post-natal day 2 (P2) fibroblast heterogeneity is fully established with the presence of the defined layers of the dermis (Figure 1a)(Driskell et al., 2013). Skin maturation occurs after P2 with the formation of the AP and the completion of the first hair follicle cycle, which results in the loss of a defined papillary fibroblast layer (Figure 1b) (Driskell et al., 2013; Rognoni et al., 2016; Salzer et al., 2018). We have previously shown that papillary fibroblasts are the primary source of de-novo dermal papilla during skin development, which are required for hair formation (Driskell et al., 2013). Furthermore, it has been suggested that adult murine skin form scars due to the lack of a defined papillary layer (Driskell et al., 2013; Driskell and Watt, 2015). Consequently, expanding this layer in adult skin might support skin regeneration in adult mice.

The use of scRNA-seq in the murine skin has established useful cell atlases of the skin during development and homeostasis (Guerrero-Juarez et al., 2019; Haensel et al., 2020; Joost et al., 2020; Joost et al., 2018; Joost et al., 2016; Mok et al., 2019). In addition, scRNA-seq studies investigating wound healing have so far focused on comparing scarring, non-scarring, or regenerating conditions (Guerrero-Juarez et al., 2019; Haensel et al., 2020; Joost et al., 2018). These studies have helped to identified key markers of the newly discovered skin fascia, such as Gpx3, which recently has been shown to contribute to scar formation (Correa-Gallegos et al., 2019; Grachtchouk et al., 2000; Joost et al., 2020). These, scRNA-seq studies have revealed that transgenic activation of the Shh pathway in the alpha-smooth-actin cells in scarring wounds, which includes pericytes, blood vessels, and myofibroblasts, can support small non-functional hair regeneration (Lim et al., 2018). However, activation of Shh pathway in dermal fibroblasts is associated with malignant phenotypes and will perturb development and homeostasis such that it may not be a safe target to support human skin regeneration clinically (Fan et al., 1997; Grachtchouk et al., 2000; Oro et al., 1997; Sun et al., 2020). Altogether, these findings suggest that an overall comparison of developing, homeostatic, scarring, and regenerating skin conditions will yield important discoveries for molecular factors that can safely support skin regeneration without harmful side effects.

The Wnt signaling pathway is involved in regulating development, wound healing, disease and cancer (Nusse and Clevers, 2017). Studies that activated Wnt and β-catenin in skin have led to important discoveries for wound healing but have produced contrasting results in the context of fibroblast biology and hair follicle formation (Chen et al., 2012; Enshell-Seijffers et al., 2010; Hamburg-Shields et al., 2015; Lim et al., 2018; Rognoni et al., 2016). Wnt is a secreted protein that activates a cascade of events that stabilizes nuclear β-catenin, which operates as a powerful co-transcription factor of gene expression. Importantly, β-catenin along cannot activate the expression of Wnt target genes without co-transcription factors. There are four Wnt co-transcription factors Tcf1 (Tcf7), Lef1, Tcf3 (Tcf7l2), and Tcf4 (Tcf7l2). These co-transcription factors modulate the functional outcome of Wnt signaling by binding to different target genes (Adam et al., 2018; Nguyen et al., 2009; Yu et al., 2012). In the context of wound healing and regeneration, it is not known how differential expression of Tcf/Lef can modulate fibroblast activity via Wnt signaling.

Since it has been shown that embryonic and neonatal skin have the potential to regenerate hair follicles upon wounding (Hu et al., 2018; Rognoni et al., 2016), we set out to identify the cell types and molecular factors that define this ability in order to transfer this regenerative potential to adult tissue. Our work has identified Lef1, as the factor in fibroblast of developing skin, that can transform adult tissue to regenerate without harmful phenotypes.

### Developing papillary fibroblasts are a transient cell population that supports hair follicle regeneration in wounds

We and others have previously shown that there are differences between neonatal and adult skin in terms of their cellular and biological properties (Ge et al., 2020; Rognoni et al., 2016; Salzer et al., 2018). These differences influence how skin can repair or if it can regenerate hair follicles in a wound. We showed that neonatal Developing skin at P2 (Figure 1a) can regenerate small hair follicles 7 days after wounding (7dpw) (Figure 1c), while adult Homeostatic skin at P21 (Figure 1b) heals by scarring 7dpw (Figure 1d). Consequently, we hypothesized that Developing or Regenerating skin contains cell types that are distinct and will support hair follicle reformation in wounds. In order to identify the cell types and molecular pathways that define the ability of Developing skin to regenerate we performed a comparison of unsorted cells from 4 conditions using scRNA-seq (Figure 1e). A total of 66,407 cells from 12 libraries (n=3 for each condition) met the preprocessing threshold and were used for downstream analysis. Each condition was represented as different colors (Figure 1a). UMAP plotting of the 66,407 sequenced cells revealed distinct clusters (Figure 1f-g). To identify cell clusters, we used Louvain modulating analysis (Joost et al., 2016; Wolf et al., 2018), which resulted in 18 distinct clusters (Figure 1g). We assigned 7 main classes of cells to the clusters. Keratinocytes represented the largest portion of cell types in the analysis (blue clusters - 0,1,2,5,8,9,11,12,17) (Figure 1h). We identified three distinct fibroblast clusters (green clusters −3,4,16). These clusters expressed Twist2/Dermo1, Pdgfra, and En1 (Figure 1h). We also identified other clusters based on their distinct gene expression profile (Figure 1h), such as immune cells (cluster 6), vasculature (cluster 7, 15), Schwann cells (cluster 10), melanocytes (cluster 13), and a pericyte population (cluster 14).

To identify a unique cell type that supports the ability to regenerate hair follicles in Developing skin we quantified the number of cells for each condition within a cluster (Figure 1i). Unexpectedly, there was no cluster defined by the Regenerating condition. Regenerating (green) and Scarring (red) conditions were found in similar proportions amongst Keratinocytes, lymphocytes/blood, melanocytes, Schwann cells, blood vessels and fibroblasts (Figure 1i). However, there were two clusters that were distinct based on the representation of cell types from a specific condition. Cluster 16 consisted of 99.9% of fibroblasts from Homeostatic skin, while Cluster 3 consisted of 67.8% of Developing fibroblasts (Figure 1i). In addition, cluster 4 consisted of 25.7% of Regenerating condition, 57.6% of the Scarring conditions. Since fibroblasts are known to play a critical (crucial) role in regulating hair follicle reformation in wounds by differentiating into de-novo DP (Driskell et al., 2013; Plikus et al., 2017; Rognoni et al., 2016), we chose to further analyze clusters 3, 4, and 16.

We subset the fibroblast clusters 3,4,16 and re-clustered them by performing an integration analysis (Polanski et al., 2020). This allowed us to test if different fibroblast subtypes, such as the papillary and reticular/hypodermal/fascia lineages, would be represented uniquely within a condition (Figure 1j-l). UMAP plotting of the integrated analysis and Louvain clustering revealed 10 distinct clusters (Figure 1k). The largest cluster (cluster 0) expressed markers of the papillary fibroblast lineage (Figure 1e-g). Four clusters (clusters 1,2,3,4) distinctly expressed markers of the reticular/hypodermal/fascia layers of the dermis (Figure 1k; Supp Fig 1a) (Correa-Gallegos et al., 2019; Joost et al., 2018). Three clusters (clusters 6,7,8) expressed markers of the dermal papilla and dermal sheath (Figure 1e-g). One cluster was distinctly represented by alpha-smooth-muscle actin indicating a myofibroblast subtype (cluster 9).

To identify the condition that is represented within specific fibroblast subtypes, we quantified the percentage of cells from each condition within the sub-clusters (Figure 1l). Our results revealed that four clusters were uniquely represented by four conditions. The myofibroblast sub-cluster (cluster 9) contained 69% of cells from the Regenerating condition, while a hypodermal fascia cluster (cluster 1) contained 92.2% of cells from the Scarring condition. Cluster 4, identified by the high expression of Cochlin (Coch), an inner ear ECM protein, contained 80.8% of cells from the Homeostasis condition and a DP/DS sub-clusters (cluster 6) contained 95.0% of cells from the Homeostasis condition. The papillary fibroblast cluster (cluster 0) contained 98.1% of cells from the Developing skin condition. The DP sub-cluster (cluster 7) consisted of 56.5% cells from the Developing condition and 37.1% cells from the Regenerating condition. These results indicate that there is a distinct separation between fibroblast sub-types that support regeneration (papillary fibroblast) or scarring (reticular/hypodermal/fascia) based on conditions.

In homeostatic skin DP and DS are critical for hair follicle growth and cycling. However, they do not migrate into wounds to support hair follicle reformation (Johnston et al., 2013; Kaushal et al., 2015). Myofibroblasts migrate into wounds but have not been shown to differentiate into dermal papilla during hair follicle regeneration and is a major characteristic of scarring (Lim et al., 2018; Plikus et al., 2017). Furthermore, reticular and fascia fibroblasts contribute to scar formation but not differentiate into dermal papilla (Correa-Gallegos et al., 2019; Driskell et al., 2013). Since our goal was to identify a cell type that uniquely represented the ability to support hair follicle regeneration in wounds, we focused on cluster 0 which are papillary fibroblasts represented by the Regenerating condition. Papillary fibroblasts migrate into wounds and support the reformation of hair follicles by differentiating into dermal papilla (Driskell et al., 2013; Rognoni et al., 2016). We conclude that the papillary fibroblasts found only in the Developing condition are unique and might harbor the potential to support hair follicle regeneration in wounds.

### Lef1 drives the papillary lineage trajectory in Developing fibroblasts

We have previously identified molecular markers of papillary, reticular and hypodermal fibroblasts in Developing P2 skin (Driskell et al., 2013). P2 developing skin is a critical time point to study the regenerative properties that can be transferrable to adult tissue. Consequently, we have generated a web-resource for this time point (https://skinregeneration.org/). Dpp4/CD26 marks the papillary fibroblast region with the highest expression with detectable signal in the reticular layer, which expresses high levels of Dlk1/Pref1 (Figure 2a-b). The hypodermis is co-labeled with Dlk1/Pref1 and Ly6a/Sca1 expression (Figure 2b-c). We subset and re-clustered the Developing P2 fibroblasts to generate a UMAP to produce a cell atlas in the context of previously defined markers of fibroblast heterogeneity (Figure 2d). Importantly, we found that PDGFRa, a pan fibroblast marker, was expressed in all populations (Figure 2d) (Collins et al., 2011). Dlk1/Pref1 and Ly6a/Sca1 labeled clusters above a population that expressed the newly identified fascia marker Gpx3 (Figure 2d). Dpp4/CD26 was detected above the Dlk1/Pref1 clusters. Dkk1, a newly identified papillary marker (Rognoni et al., 2016), was expressed in an adjacent population to the Dpp4/CD26 population. MKi67 marked cells undergoing division, but importantly overlapped with Dpp4/CD26 and Dkk1 expression, suggesting that this cluster represented dividing papillary fibroblasts. Acan, a newly identified DS marker (Heitman et al., 2020), was detected in a small population adjacent to the cluster which expressed the DP marker Corin (Enshell-Seijffers et al., 2008). We conclude that our scRNA-seq analysis of Developing fibroblasts supports previously identified markers defining fibroblast lineages but expose the existence of two papillary fibroblast populations, suggesting additional heterogeneity to be investigated in future studies.

**Figure 2:**
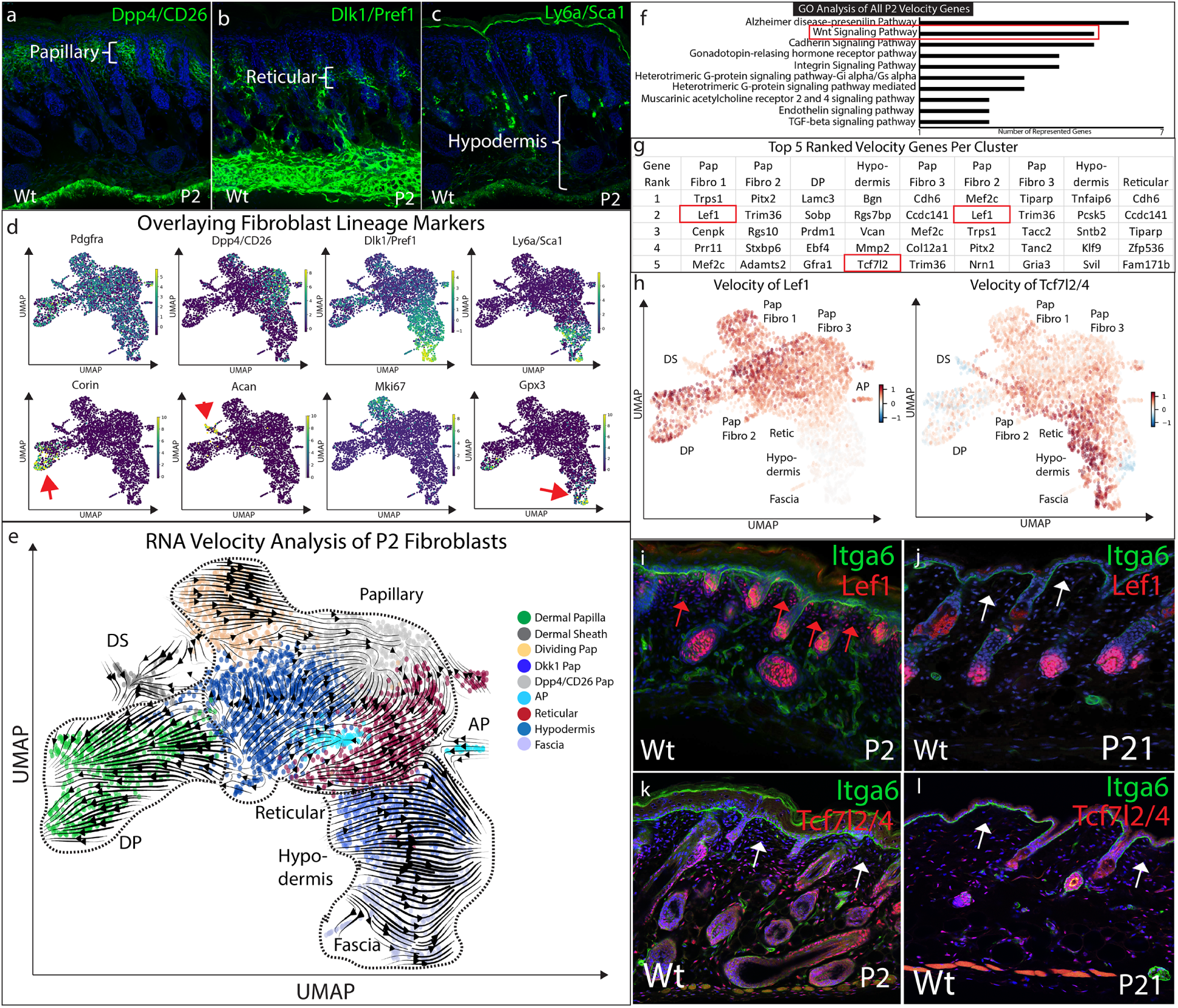
Papillary fibroblast lineage trajectory is defined by Lef1 in RNA Velocity analysis. (a-c) Immunostaining Development (P2) skin for Papillary (a) (Dpp4/CD26), Reticular/Hypodermal (b) (Dlk1/Pref1), and Hypodermal makers (c) (Ly6a/Sca1). (d) Overlaying fibroblast lineage markers on UMAP projections of Developing fibroblasts. (e) RNA Velocity analysis over layed on UMAP projection of clustered fibroblasts. Clusters are colored coded and labeled. (f) GO Analysis of 195 driver genes of RNA Velocity for Developing fibroblasts. (g) Top 5 Velocity driver genes of Developing fibroblast clusters. (h) Overlaying Lef1 and Tcf7l2/4 velocity on UMAP projections of Developmental fibroblast. (i-l) Immunostaining Developing (P2) and Homeostatic (P21) time points for Lef1 and Tcf7l2/4 expression and counterstained with Itga6. Red arrows indicate areas of papillary dermis expressing Lef1. White arrows indicate papillary dermal region.

To identify factors that define the papillary fibroblast clusters we performed an RNA Velocity analysis on Developing fibroblasts (Figure 2e) (Methods: Github weblink). RNA Velocity is a function of scRNA-seq that computationally compares the ratio of spliced/unspliced mRNA sequencing data, within a cell, to establish an estimated trajectory of differentiation, projected as arrows on a UMAP plot (Figure 2e) (La Manno et al., 2018). The arrows are defined by a list of driver genes (Supp Table 1). Interestingly, the estimated trajectories of the clusters support our previous work indicating that cells that express Ly6a/Sca1 do not differentiate into the papillary lineage (Driskell et al., 2013). We probed the list of Velocity driver genes (Supp Table 2) using GO Pathway analysis (Figure 2f). The Wnt signaling pathway was one of the most significant pathways represented. RNA Velocity analysis also ranks the top 5 drivers for each cluster, which revealed two canonical Wnt transcription factors with opposing trajectories (Figure 2g-h). Lef1, driving trajectories of 2 papillary clusters and Tcf4 driving the hypodermal cluster. Importantly, Lef1 is an early DP marker capable of enhancing human hair follicle growth in vitro, while Tcf4 is associated with fibrotic scars (Mok et al., 2019) (https://doi.org/10.1101/2020.01.05.895334). We projected the velocity of Lef1 and Tcf4 as gradients of color on UMPAs of Developing fibroblasts (Figure 2h) to visualize the direction of their trajectories. The velocity of Lef1 was positive in the direction of all papillary clusters including the dermal papilla and dermal sheath. Tcf4 velocity was positive toward the reticular/hypodermal/fascia clusters. To validate Lef1 expression in papillary fibroblasts, we immunostained Developing (P2) and Homeostatic (P21) tissue samples (Figure 2i-l). We found Lef1 expression in the papillary region of Developing skin, which was lost in adult dermis and correlates with the loss of regenerative ability in Homeostatic skin. Likewise, we confirmed that Tcf4 was detected in the reticular/hypodermis throughout Developing skin and in Homeostasis but not in papillary fibroblasts. Interestingly, it has been previously shown that the response to Wnt signals could be specified by the different Tcf factors expressed (Adam et al., 2018). Since Lef1 has previously been defined as an early DP marker (Mok et al., 2019) we hypothesize that Lef1 marks papillary fibroblasts capable of supporting hair follicle regeneration in wounds.

### Neonatal skin regeneration requires Lef1 expression in fibroblasts

Based on our scRNA-seq findings, we hypothesized that Lef1 defines a neonatal papillary fibroblast that supports skin regeneration during neonatal wound healing. To test if Lef1 in fibroblasts is required to support neonatal regeneration, we produced a tissue specific knockout model. We utilized the fibroblast specific promoter Dermo1/Twist2 to drive Cre expression, bred with a mouse line with flox sites flanking Exon 7 and 8 of the Lef1 locus (Yu et al., 2012). We called this mouse line DermLef1KO (Figure 3a). DermLef1KO mice were viable and fertile with small shifts in hair follicle phenotypes that resulted in less dense fur (Manuscript in Prep). We confirmed dermal Lef1 ablation from the papillary fibroblast at P2 and from adult DP by immunostaining (Figure 3b-e). We also performed 2mm wounds on P2 DermLef1KO and wild type littermates harvesting at 7dpw (Figure 3f). Our analyses revealed that Lef1 was expressed in the de-novo DP and regenerating hair follicle buds of wild type mice (Figure 3g,i), but that wounded DermLef1KO mice lacked regeneration (Figure 3h,j,k). We conclude that Lef1 expression in neonatal fibroblasts is required for hair follicle regeneration in wounds.

**Figure 3:**
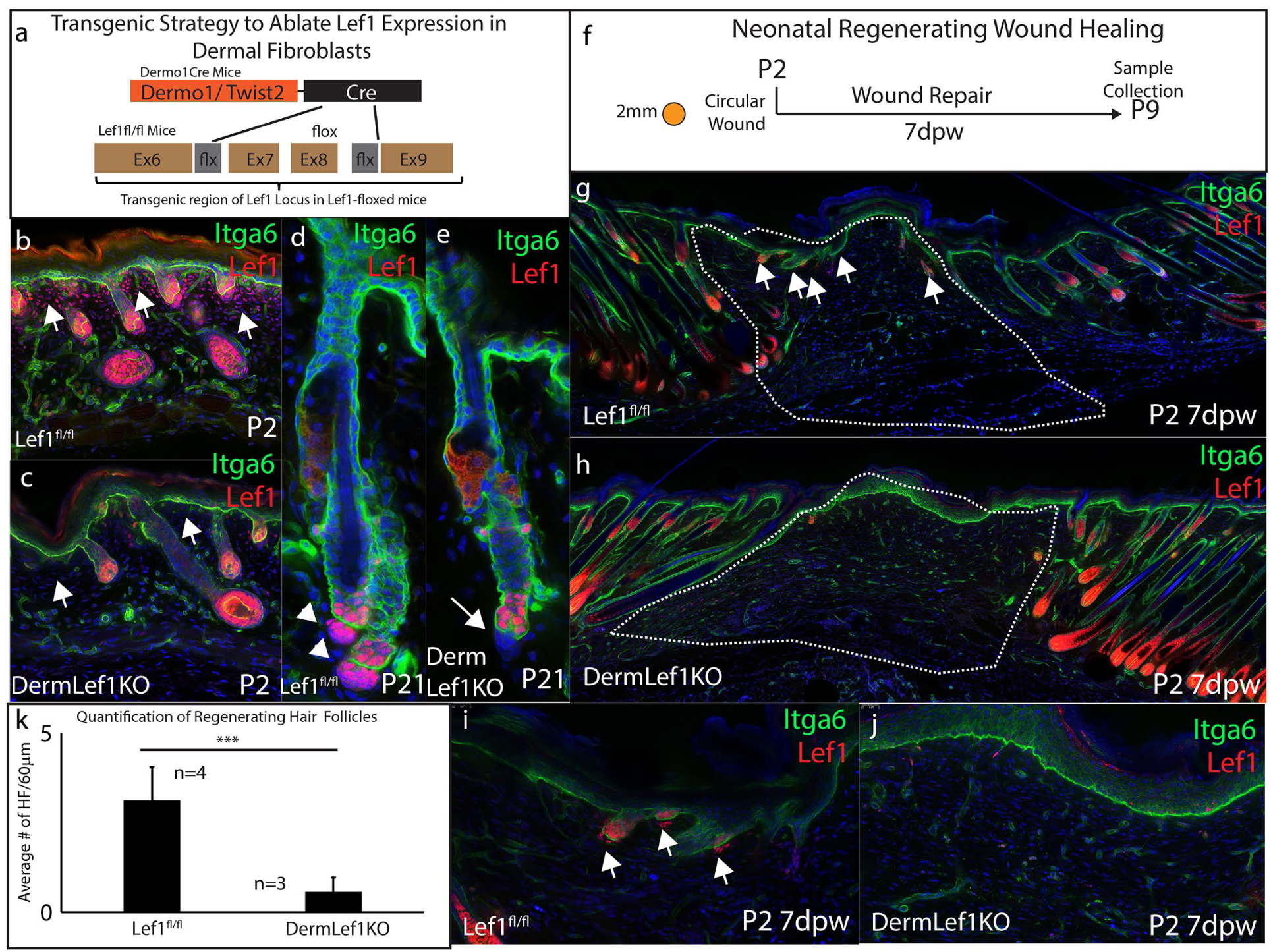
Dermal Lef1 is required to support hair follicle regeneration in neonatal wounds. (a) Schematic representation of transgenic strategy to ablate Lef1 expression in dermal fibroblasts. (b-e) Verifying dermal Lef1 ablation by immunostaining. White arrows indicate papillary regions (b-c) or dermal papilla (d-e). (f) Schematic describing 2mm circular wounds and harvest times to test if dermal Lef1 is required for regeneration. (g-i) Immunostaining of P2 7 days post wound (7dpw) skin for Itga6 and Lef1. Wound beds are highlighted by white outline. White arrows indicate regenerating hair follicles. (k) Quantification of regenerating hair follicles in wounds beds of n=4 Lef1fl/fl (wt) and n=3 DermLef1KO mice. p < 0.003

### Dermal Lef1 expression primes adult skin to enhance regeneration in large wounds

Our scRNA-seq data, together with tissue specific knockout, suggests that skin can be primed to regenerate during development and homeostasis. We hypothesize that constitutively active Lef1 expression in dermal fibroblasts will prime skin to support regeneration. To induce Lef1 expression in fibroblasts, we utilized a previously published transgenic mouse model Lef1KI (Lynch et al., 2016) bred with Dermo1/Twist2-Cre mouse line called the DermLef1KI line (Figure 4a). We verified that Lef1 was expressed in all dermal fibroblast compartments in DermLef1KI mice during development and in adult homeostatic skin conditions (Figure 4b-e). Importantly, the only phenotype detected was the early entry of the hair follicle cycling into anagen at P21 in DermLef1KI mice (Figure 2d-e). In addition, DermLef1KI mice have similar life spans as wild type mice without any adverse effects (Supp Fig 3). Our results reveal that induced expression of Lef1 in fibroblasts does not negatively affect the development and homeostasis of murine skin.

**Figure 4:**
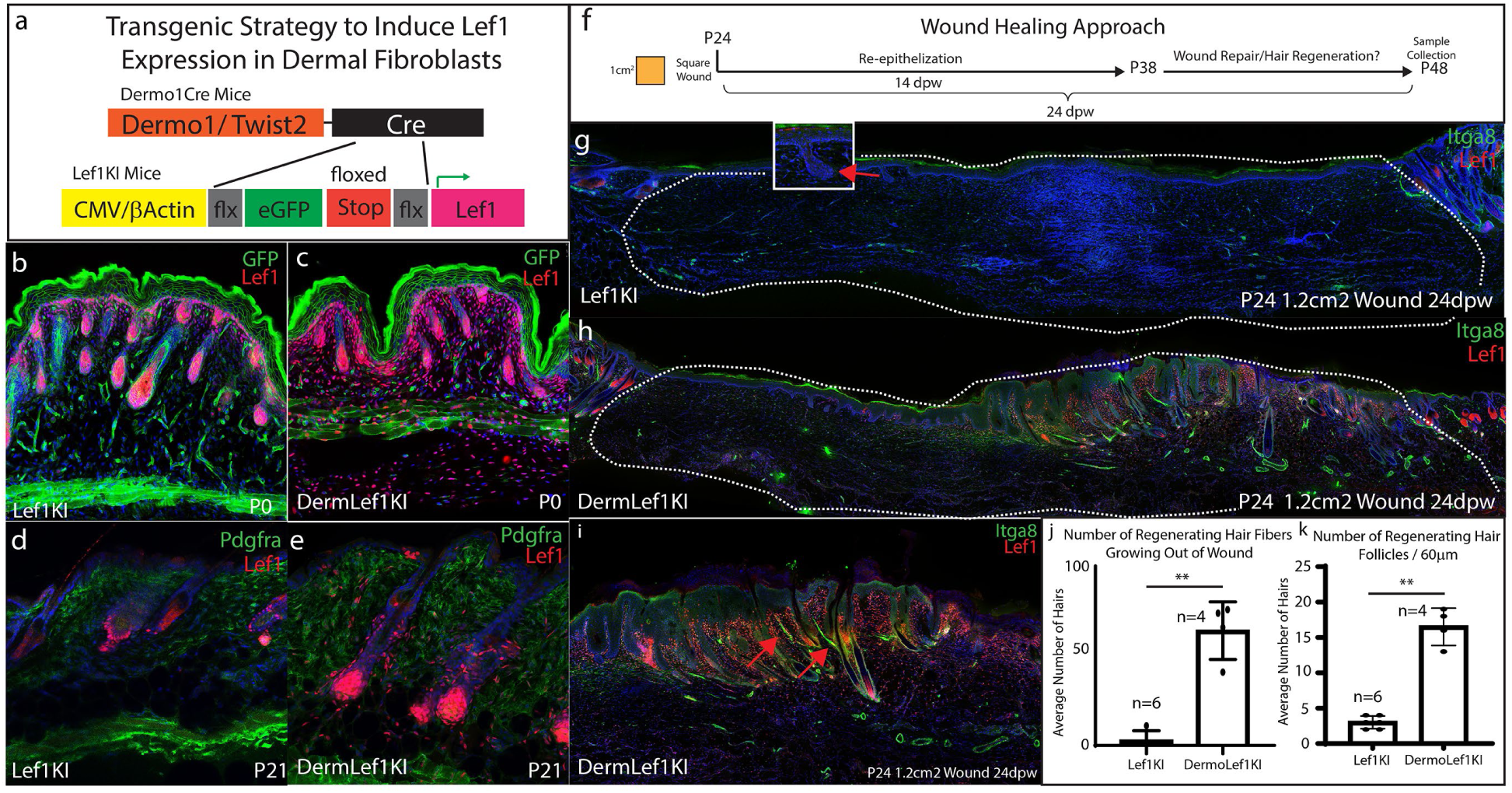
Lef1 expression in dermal fibroblast enhances skin regeneration in large wounds. (a) Schematic representation of transgenic strategy to ectopically express Lef1 in fibroblasts during development, homeostasis and wound healing. (b-c) Immunostaining for Lef1 and GFP in P0 Lef1KI and DermLef1KI (Dermo1Cre+/Lef1KI+) mice. (d-e) Immunostaining for Lef1 and PDGFRa in P21 Lef1KI and DermLef1KI mice. (f) Schematic representation of wound healing approach. (g-k) Immunostaining tissue for Lef1 and Itga8 (arrector pili) from the wounds of Lef1KI and DermLef1KI mice. Regenerating hair follicles in Lef1KI mice can be found by red arrows. Wound beds are marked by red squares. High resolution area of regenerating hair follicles in DermLef1Ki mice (i). (j) Quantification of regenerating hair follicles growing out of the wound beds of Lef1KI n=6 and DermLef1KI n=4 mice p < 0.004. (k) Quantification of regenerating hair follicles detected in 60 μm sections of wound beds. p < 0.001

Adult skin normally does not regenerate but has the capacity to reform non-functional hair follicles (small and lack arrector pili) in wounds 1cm^2^ or larger.(Ito et al., 2007) To test if overexpression of Lef1 in adult dermal fibroblasts enhances hair regeneration in adult skin, we wounded wild type (Lef1KI) and DermLef1KI mice at P24 with 1.2cm^2^ wounds and harvested at 24dpw (Figure 4f). Our analysis comparing Lef1KI and DermLef1KI wound beds revealed enhanced hair regeneration (Figure 4f-k). There were 5 times more regenerating hair follicles on average in DermLef1KI wound beds compared to wild type wounds (Figure 4j-k). The non-functional hair follicles that typically regenerate in large adult wounds are small and lack the arrector pili, a smooth muscle that makes hair stand up when contracted. Strikingly, the hair follicles regenerating in the DermLef1KI wound beds contained arrector pili (Figure 4i) and were larger than hair follicles found in wild type wounds. This extensive ability to regenerate has only been reported to occur in the African Spiny Mouse (Acomys) model.(Jiang et al., 2019; Seifert et al., 2012) We conclude that Lef1 expression in dermal fibroblasts primes skin to enhances regeneration.

### Dermal Lef1 transforms scarring wounds to be regenerative

Wound healing studies utilize a standardized approach to take into account the hair follicle cycle, which influences the wound healing process.(Ansell et al., 2011) Consequently, the bulk of wound healing and regeneration studies are performed at 3 weeks of age (∼P21) or at 7-8 weeks of age (∼P57) (Gay et al., 2013; Guerrero-Juarez et al., 2019; Ito et al., 2007; Lim et al., 2018; Plikus et al., 2017; Plikus et al., 2008). In addition, a standard size wound of roughly 8mm in diameter or smaller is used, which heal by scarring (Driskell et al., 2013; Lim et al., 2018; Rognoni et al., 2016). Since Lef1KI mice showed enhanced regenerative capacity in large wounds, we explored the regeneration potential of DermLef1KI skin, in wounds that normally scar. To determine if the hair follicle cycle influences skin regeneration we performed wounds at different time points. P24 which undergoes a hair follicle cycle during wound healing. P40 which heals during the resting phase of the hair cycle. And P90 which is the beginning of the un-synchronized hair follicle cycling that occurs in adult mice (Figure 5a). Scars formed in wounds performed at P24 in both WT and DermLef1KI mice (Figure 5b-c,l). However, hair follicle regeneration with arrector pili occurred in wounds performed at P40 and P90 in DermLef1KI mice (Figure 5d-k, l, Supp Fig 4) (Figure 4h-k). We conclude that Lef1 expression in fibroblasts induces skin to support hair follicle regeneration in wounds that heal outside of the hair follicle cycle.

**Figure 5:**
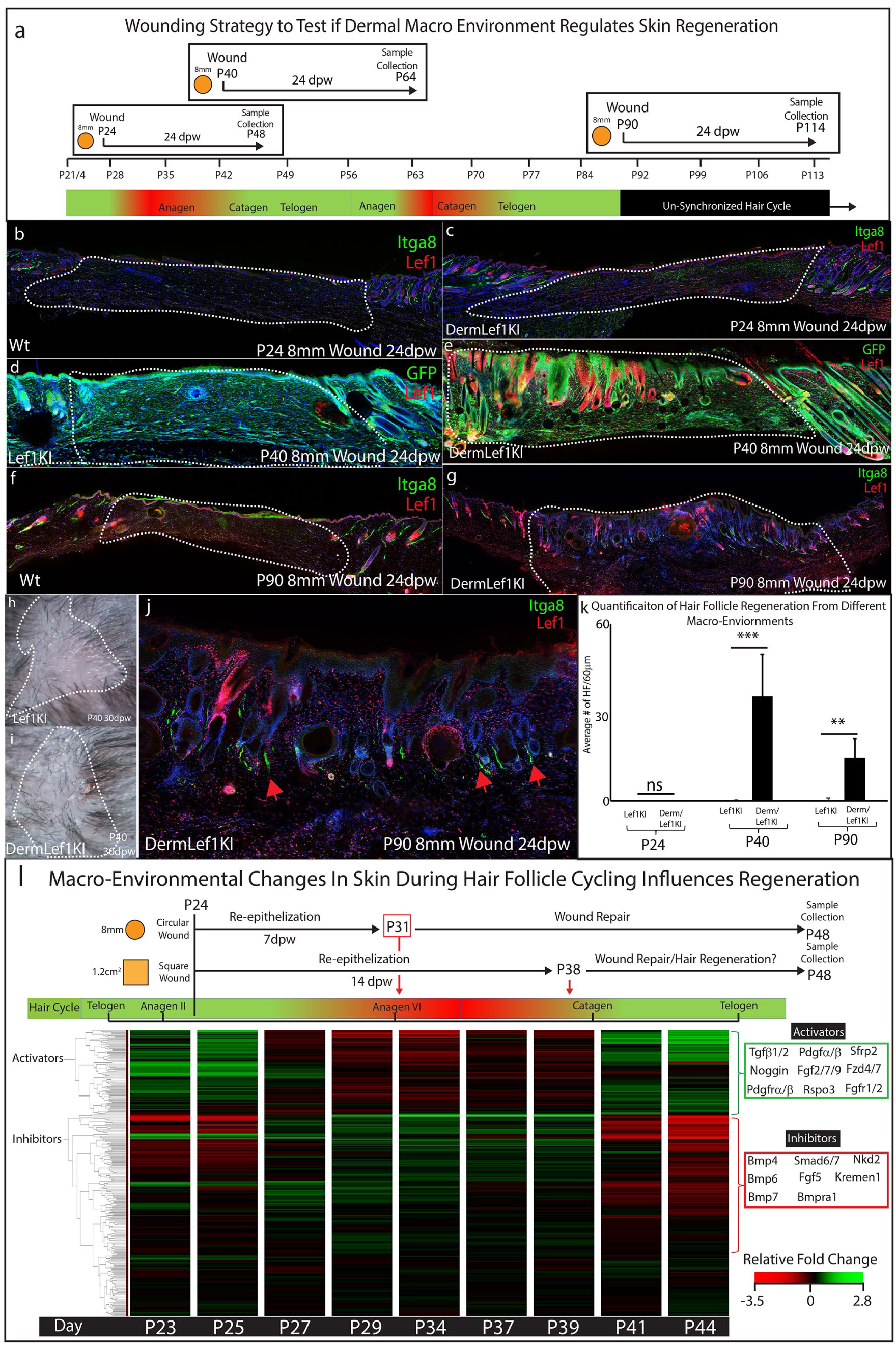
Lef1 expression in dermal fibroblasts enhances skin regeneration in permissive macro environments. (a) Schematic of the differential wound healing analysis from different time points of the hair follicle cycle in murine skin. (b-g,j) Immunostaining analysis of wounds 24dpw from P24 (b-d), P40 (d-e), and P90 (f-g,j). Sections were stained for either Itga8 and Lef1 (b-c,f-g,j) or GFP and Lef1 (d-e). Itga8 labels arrector pili. (h-i) Macroscopic analysis of wound beds from Wt (Lef1KI) and DermLef1KI mice. Hair follicles could be seen growing out of the wound bed of DermLef1KI mice. (k) Quantification of the average number of regenerating hair follicles detected in wounds beds of 60 μm sections from P24 (ns) 24dpw (p < 0.0009), P40 24dpw, and P90 24dpw (p < 0.007). (l) Schematic representation of different wound healing approaches above a heatmap of microarray data (Lin et al., 2009) from key time points during hair follicle cycling. Heatmap is generated from a list of Activator and Inhibitor genes reported to regulate the progression of the hair follicle cycle (Wang et al., 2017).

Our wound healing studies mirror previous work suggesting that the size of a wound can dictate the potential of skin to regenerate instead of scar (Ito et al., 2007). To explain ‘why size matters’ we investigated and mapped the timing of wound closure and regeneration in the context of molecular signals within the dermal macro-environment that are dynamically changing throughout the hair follicle cycle (Plikus et al., 2008; Wang et al., 2017). Previous studies have defined the activators and inhibitors that regulate hair follicle cycling during this time (Wang et al., 2017) (Supp Table 3). Activators include the Wnt signaling pathway, Noggin, Tgfb, and fibroblast markers such as PDGFRa. Inhibitors include the BMP signaling pathway. These activators and inhibitors have been previously shown to modulate Lef1 activity (Jamora et al., 2003). To explain why DermLef1KI mice scar at P24, we generated a diagram of wound re-epithelization and healing time points comparing 8mm circular wounds to 1.2cm^2^ wound in the context of gene expression for inhibitors and activators at all time points during the hair follicle cycle (Figure 5l). 8mm circular wounds re-epithelialize 7dpw (Driskell et al., 2013) in a macro-environment with high levels of Lef1 inhibitors, which are not inductive to hair follicle formation (Figure 5l). In contrast, large 1.2cm^2^ wounds re-epithelialize during the time point of the hair cycle that has high activators and low inhibitors (Figure 5l). We conclude that the size of a wound dictates the time point as to when a wound closure is associated with inhibitor or inductive regenerative signals.

## Discussion

It is well established that embryonic and neonatal skin has the potential to heal in a scar-less manner (Walmsley et al., 2015a). However, the cell types and signals that can transfer this ability to adult skin have not been identified. In this study we identify Developing papillary fibroblasts as a transient cell population that is defined by Lef1 expression. Inducing Lef1 expression in adult fibroblasts will prime skin to support enhanced hair follicle regeneration. Interestingly, ectopic Lef1 expression in fibroblasts did not result in any adverse phenotypes with phenotypically normal mice (Supp Fig 3).

scRNA-seq has provided an unprecedented insight in the cell types and molecular pathways that are represented within regenerating and scarring wounds (Guerrero-Juarez et al., 2019; Haensel et al., 2020; Joost et al., 2018). For example, comparisons of these different repairing conditions have identified the activation of the Shh pathway as a key pathway to support hair follicle regeneration in wounds of different sizes (Lim et al., 2018). However, the activation of ectopic Shh in either the epidermis or dermis is associated with unwanted phenotypes and cancer (Fan et al., 1997; Grachtchouk et al., 2000; Oro et al., 1997; Sun et al., 2020). Unexpectedly, we discovered a transient papillary fibroblast population that is defined by Lef1 expression in Developing skin instead of in Regenerating conditions. Importantly, our comparative analysis of Regenerating and Scarring wounds reveals their similarities (Figure 1i) more than their differences suggesting that the wound environment is a powerful driving force for gene expression even in a heterogeneous population of cells. Consequently, the basic mechanisms occurring during development and maintenance of neonatal tissue may hold the keys to transforming adult tissue to regenerate instead of scarring.

Studies involving the activation of the Wnt and β-catenin pathways in skin have led to important discoveries in wound healing research, but have produced contrasting results in the context of fibroblast biology and hair follicle formation (Chen et al., 2012; Hamburg-Shields et al., 2015; Ito et al., 2007; Lim et al., 2018; Mastrogiannaki et al., 2016; Rognoni et al., 2016). The activation of β-catenin in fibroblasts during wound healing has been shown to inhibit hair follicle reformation in scars (Lim et al., 2018; Rognoni et al., 2016) while deactivating Wnt signaling inhibited hair reformation (Myung et al., 2013). Moreover, persistent Wnt activity in the dermis has been recently shown to drive WIHN towards fibrotic wound healing (Gay et al., 2020). In contrast, the activation of β-catenin during development increases hair formation and size (Chen et al., 2012; Enshell-Seijffers et al., 2010). One explanation for these conflicting results is the presence of different fibroblast lineages present in either development or wounds, which express different Lef/Tcf factors to direct a Wnt/β-catenin specific response (Driskell et al., 2013; Philippeos et al., 2018; Rognoni et al., 2016). We and others have shown that Lef1 is expressed in embryonic and neonatal papillary fibroblasts, which is lost in adult fibroblasts (Driskell et al., 2013; Mastrogiannaki et al., 2016; Philippeos et al., 2018; Rognoni et al., 2016). Here we demonstrate that Lef1 is a key inductive element in developing fibroblasts that supports hair follicle formation in adult wounds. Our data indicates that Lef1 expression in fibroblasts primes a response to Wnt signals to support hair follicle neogenesis and arrector pili reformation, without negative affects to development or homeostasis, as seen in other models (Seifert et al., 2012). Our results align with studies that induced Lef1 expression in cultured human dermal papilla cells to enhance hair follicle formation in vitro (Abaci et al., 2018). In humans, at the age of 50, the quality of the papillary dermis gradually deteriorates, which correlates with decreased wound healing abilities (Haydont et al., 2019; Marcos-Garces et al., 2014). Our data suggests that activating the human papillary region to retain its identity by expression of Lef1 has the potential to enhance wound healing in humans.

## Material and Methods

### Mouse models

All mice were outbred on a C57BL6/CBA background, with male and female mice used in all experiments. The following transgenic mouse lines were used in this study, Twist2-Cre (Cat# 008712). The ROSA26-CAG-flox-GFP-STOP-flox-Lef1 and Lef1-flox-Exon7,8-flox mouse has been previously described (Lynch et al., 2016; Yu et al., 2012). Wild type mice utilized in scRNA-seq experiments were an outbred background of C57BL6/CBA mice.

### Wounding experiments

All wounding experiments were done in accordance with the guidelines from approved protocols Washington State University IACUC. Mice were wounded at post-natal day (P) 2, P24, P40 and P90 as stated in the results section. All wounding experiments were carried out with mice under anesthetization. The dorsal hair was shaved, and wounding areas were disinfected. For full-thickness circular wound, 2mm and 8mm punch biopsy and surgical scissors were used on P2 and P24/P40 mice, respectively. For large wounds, 1.44cm2 (1.2 x 1.2 cm) squares were excised using surgical scissors. Wounds were harvested at 7 dpw, 14 dpw, 24 dpw, and 30 dpw.

### Horizontal whole mount

This procedure was described in Salz et al., 2017 (Salz and Driskell, 2017). Briefly, full thickness skin samples were collected and fixed in 4% Paraformaldehyde (PFA), before being frozen in OCT compound in cryo-block. Samples were sectioned using cryostat at 60 mm thickness. Skin sections were then stained with primary antibodies at 4C overnight in PB buffer and with secondary antibodies in PB buffer for one hour. Samples were mounted on glass cover slip with glycerol as mounting medium.

### Generation of Single cell RNA-Sequencing data

Cells isolation procedure was previously described in Jensen and Driskell 2009 (Jensen et al., 2010). Single-cell cDNA libraries were prepared using the 10X Genomics Chromium Single Cell 3’ kit according to the company’s instruction. For each condition, 3 mice were used to generate 3 separate libraries, making it a total of 12 libraries across all 4 conditions. All samples were processed individually. The prepared libraries were sequenced on Illumina HiSeq 4000 (100 bp Paired-End). Demultiplexed Paired-End fastq files were aligned to mm10 reference genome using 10X Genomics CellRanger function (Cellranger version 3.0.2). The outputs of Cellranger alignment were used to create loom files by Velocyto “run10x” function (La Manno et al. 2018). Loom files are the preferred input data for scVelo package to perform RNA Velocity analysis (Bergen et al. 2019).

### Single cell RNA-Seq data preprocessing

For basic filtering of our data, we filtered out cells have expressed less than 200 genes and genes that are expressed in less than 3 cells. To ensure that the data is comparable among cells, we normalized the number of counts per cell to 10,000 reads per cell as suggested by the scanpy pipeline (Wolf et al. 2018). Data were then log-transformed for downstream analysis and visualization. We also regressed out effects of total counts per cell and the percentage of mitochondrial genes expressed, then scaled the data to unit variance with the maximum standard deviation 10.

### Single cell RNA-Seq data analysis

Neighborhood graph of cells was computed using PCA presentation (n PCs = 40, n neighbors = 10). The graph was embedded in 2 dimensions using Uniform Manifold Approximation and Projection (UMAP) as suggested by scanpy developers. Clusters of cell types were defined by Louvain method for community detection on the generated UMAP graph at resolution of 0.2. The analysis pipeline is published on the Driskell lab Github page as Jupyter Notebook (https://github.com/DriskellLab/Priming-Skin-to-Regenerate-by-Inducing-Lef1-Expression-in-Fibroblasts-).

### Microarray analysis

The microarray data were previously published (GEO GSE11186) (Lin et al., 2009). RMA normalization was used, and the bottom twentieth percentile of genes were excluded from the analysis.

### Data availability

All data presented in this manuscript are available from the authors upon request. The detail analysis of scRNA-Seq and RNA velocity can be found at https://github.com/DriskellLab/Priming-Skin-to-Regenerate-by-Inducing-Lef1-Expression-in-Fibroblasts-. We have also generated a web resource that allows for easy access to our data through a query-based display of our scRNA-Seq analysis, available at https://skinregeneration.org/. This website will be continuously updated with more dataset and analysis.

## Acknowledgements

This work was supported by WSU New Faculty Seed Grant 2428-9926. The authors would like to acknowledge Sean Thompson, Jonathan Jones, Kai Kretzschmar, and Klaas Mulder for their discussion and help. The authors would also like to acknowledge the Washington State University Franceschi Microscopy Imaging Center for the continued support in microscopic imaging, and Jared Brannan for generation of our web resource.

## Author Contributions

QP performed and designed the experiments, analyzed the data, and co-wrote the manuscript. GF, LS, GGH, BW, and IMD assisted in performing and designing experiments. RRD conceived of the project, assisted in experimental design, performed data analysis, and co-wrote the manuscript.

---

**Supplemental Figure 1:**
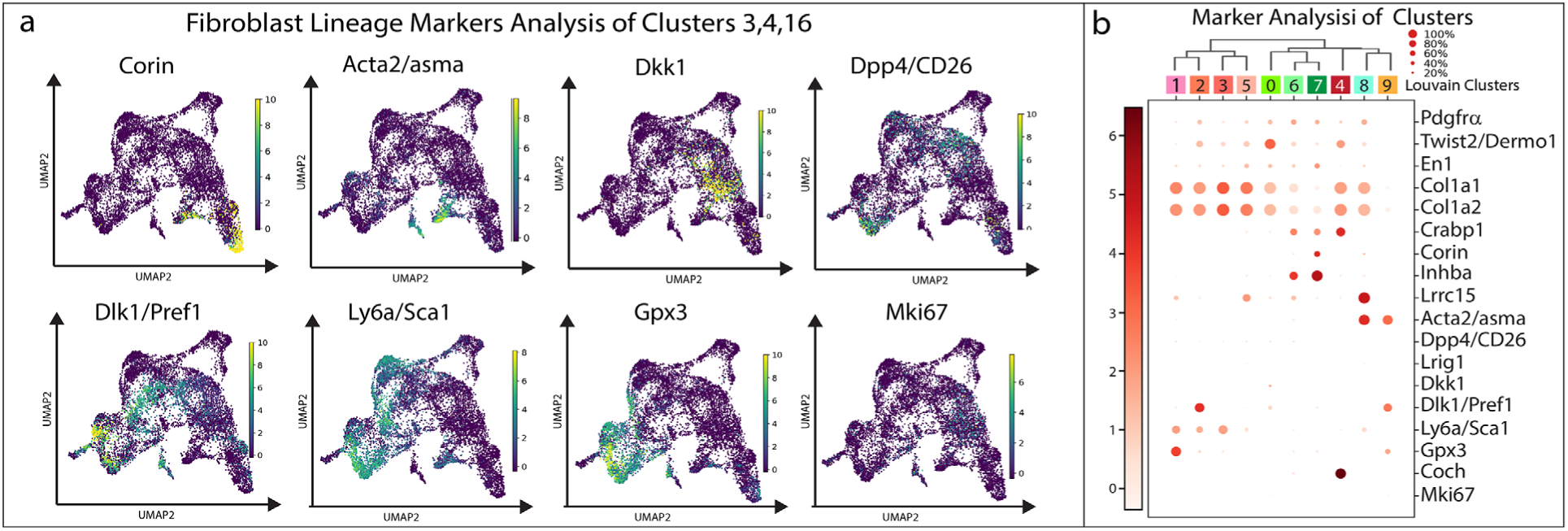
Classifying fibroblast clusters and conditions in Development, Regeneration, Scarring, and Homeostasis. (a) Over-laying gene expression for cell markers on UMAP plots. (b) Dotplot of genes based on expression in cluster.

**Supplemental Figure 2:**
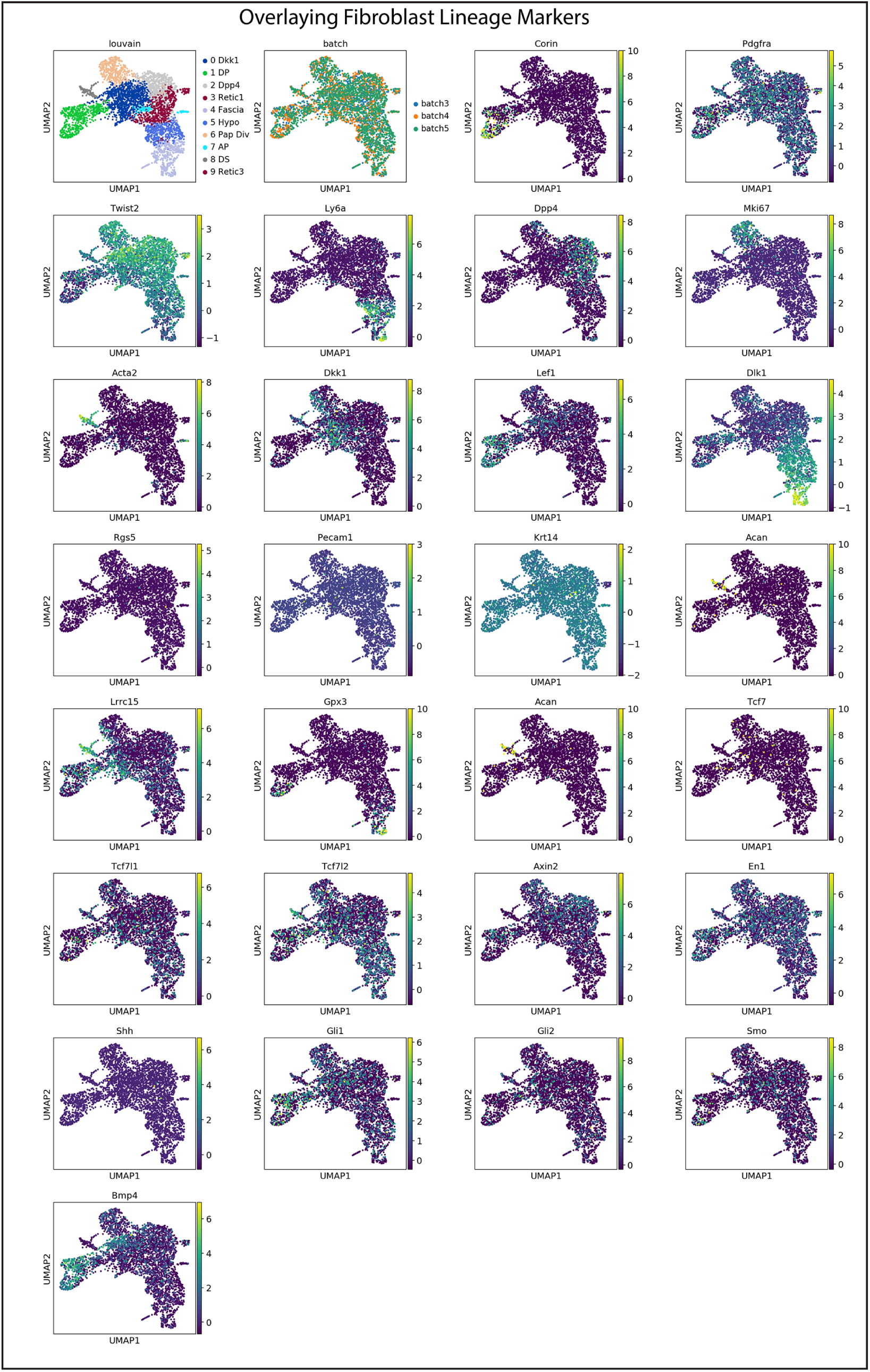
Classifying fibroblast clusters in Developing fibroblasts. (a) Over-laying gene expression for fibroblast lineage markers.

**Supplemental Figure 3:**
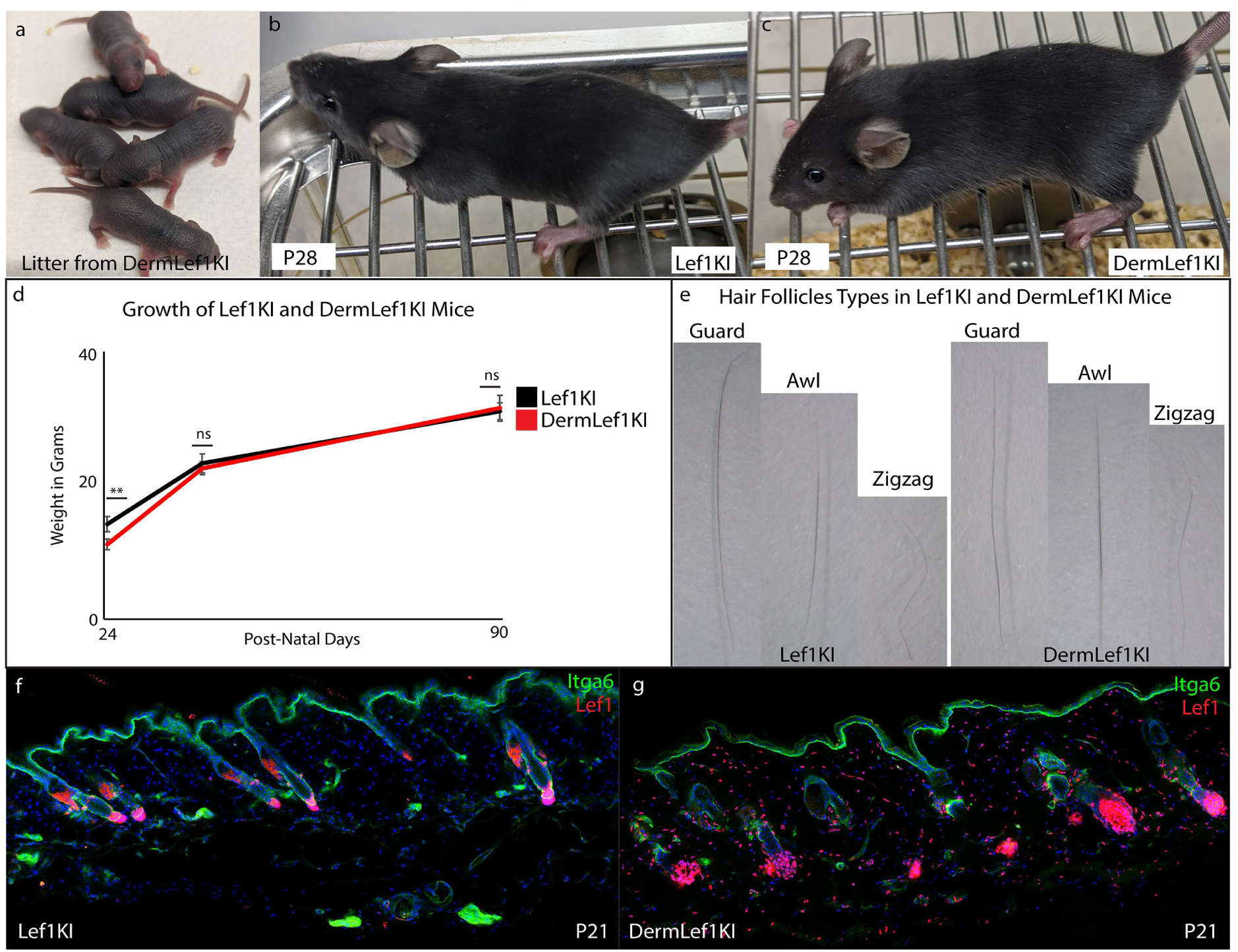
DermLef1KI mice does not show overt phenotypic variations. (a) Images of P4 litter from DermLef1KI mice revealing viability of line. (b-c) P28 of wild type (b) and DermLef1KI (c) mice. (d) The growth curve of Wt (Lef1KI) mice compared to DermLef1KI mice at P24, P40, and P90. P24 p < 0.005. (e) Macroscopic images of different hair follicle types from Wt (Lef1KI) and DermLef1KI hair follicles. (f-g) Immunostaining analysis of sections from P21 skin from Wt (Lef1KI) and DermLef1KI mice stained for Itga6 and Lef1.

**Supplemental Figure 4:**
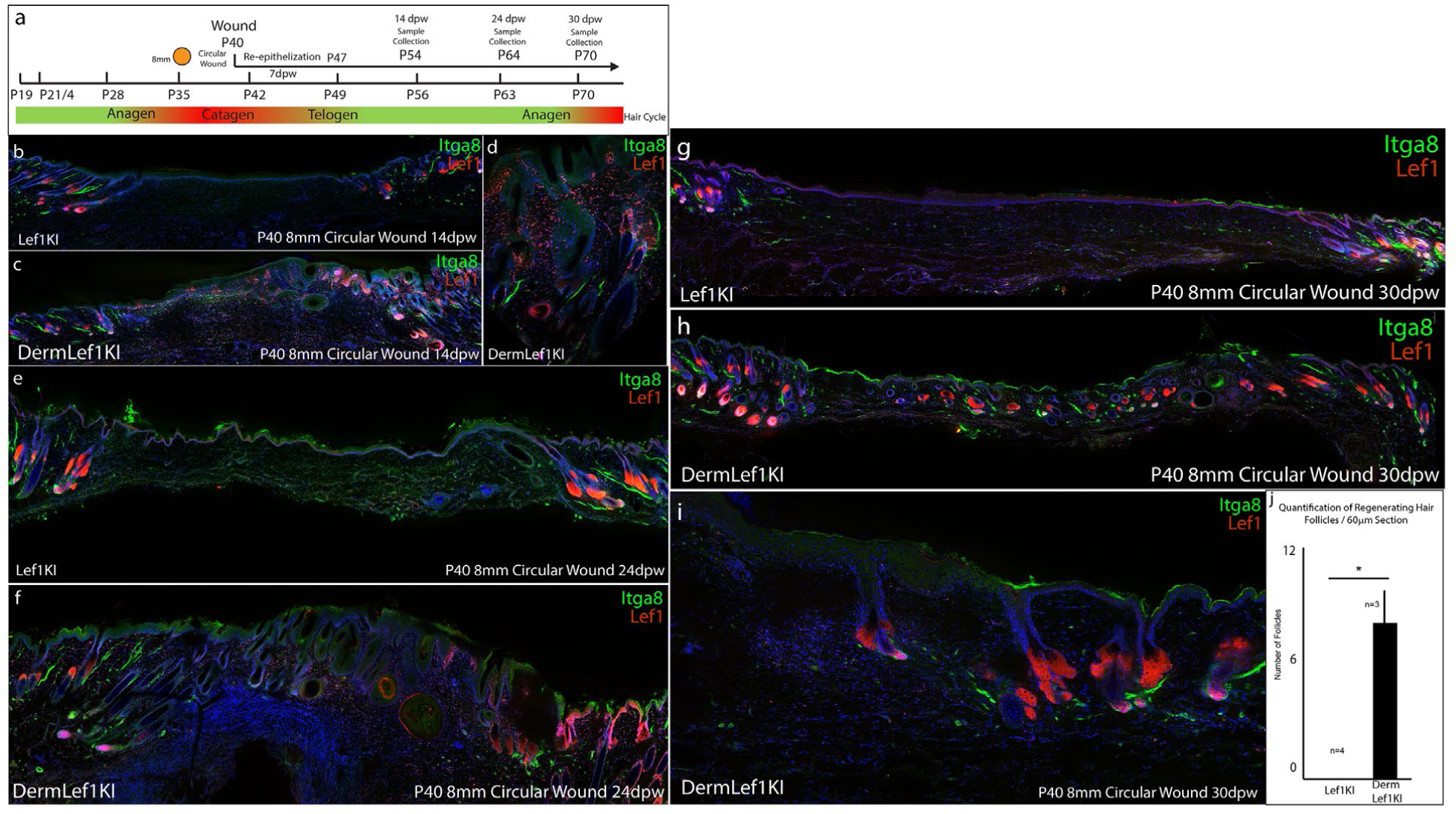
Analysis of the regenerative potential of DermLef1KI wounds at P40. (a) Schematic describing the wounding and tissue collection strategy. (b-j) Immunostaining analysis of wild type (Lef1KI) and DermLef1KI mice wounded at P40 and collected at 14 dpw (b-d), 24dpw (e-f), and 30dpw (g-i). All sections were immunostained for Itga8 and Lef1. (j) Quantification of hair follicles in 60μm sections of Wt (Lef1KI) and DermLef1KI skin 30dpw. p < 0.04

